# A New Power Law Linking the Speed to the Geometry of Tool-Tip Orientation in Teleoperation of a Robot-Assisted Surgical System

**DOI:** 10.1101/2022.03.02.482648

**Authors:** Or Zruya, Yarden Sharon, Hanna Kossowsky, Fulvio Forni, Alex Geftler, Ilana Nisky

**Affiliations:** Department of Biomedical Engineering and the Zlotowsky Center for Neuroscience, Ben-Gurion University of the Negev, Beer Sheva, Israel; Department of Orthopedic Surgery, Soroka Medical Center, Ben-Gurion University of the Negev, Beer Sheva, Israel; Department of Engineering, University of Cambridge, Cambridge CB2 1PZ, U.K.

## Abstract

Fine manipulation is important in dexterous tasks executed via teleoperation, including in robot-assisted surgery. Discovering fundamental laws of human movement can benefit the design and control of teleoperated systems, and the training of their users. These laws are formulated as motor invariants, such as the well-studied speed-curvature power law. However, while the majority of these laws characterize translational movements, fine manipulation requires controlling the orientation of objects as well. This subject has received little attention in human motor control studies. Here, we report a new power law linking the speed to the geometry in orientation control – humans rotate their hands with an angular speed that is exponentially related to the local change in the direction of rotation. We demonstrate this law in teleoperated tasks performed by surgeons using surgical robotics research platforms. Additionally, we show that the law’s parameters change slowly with the surgeons’ training, and are robust within participants across task segments and repetitions. The fact that this power law is a robust motor invariant suggests that it may be an outcome of sensorimotor control. It also opens questions about the nature of this control and how it can be harnessed for better control of human-teleoperated robotic systems.

## I. INTRODUCTION

In many teleoperated robot-assisted applications, users perform tasks requiring fine manipulations in six degrees of freedom - the translation and the rotation of objects (Fig. 1). For example, in Robotic-Assisted Minimally Invasive Surgeries (RAMIS), surgeons teleoperate robotic manipulators to control instruments inside a patient’s body with which they manipulate objects such as tissue and needle. Understanding how humans coordinate the movement of their hands can be beneficial in the design and control of teleoperated systems, and the training of their users. It can also be informative for the design of path planners and controllers for systems that operate together with human users by making these systems more human-like and thus predictable [1], [2], [3].

**Fig. 1.**
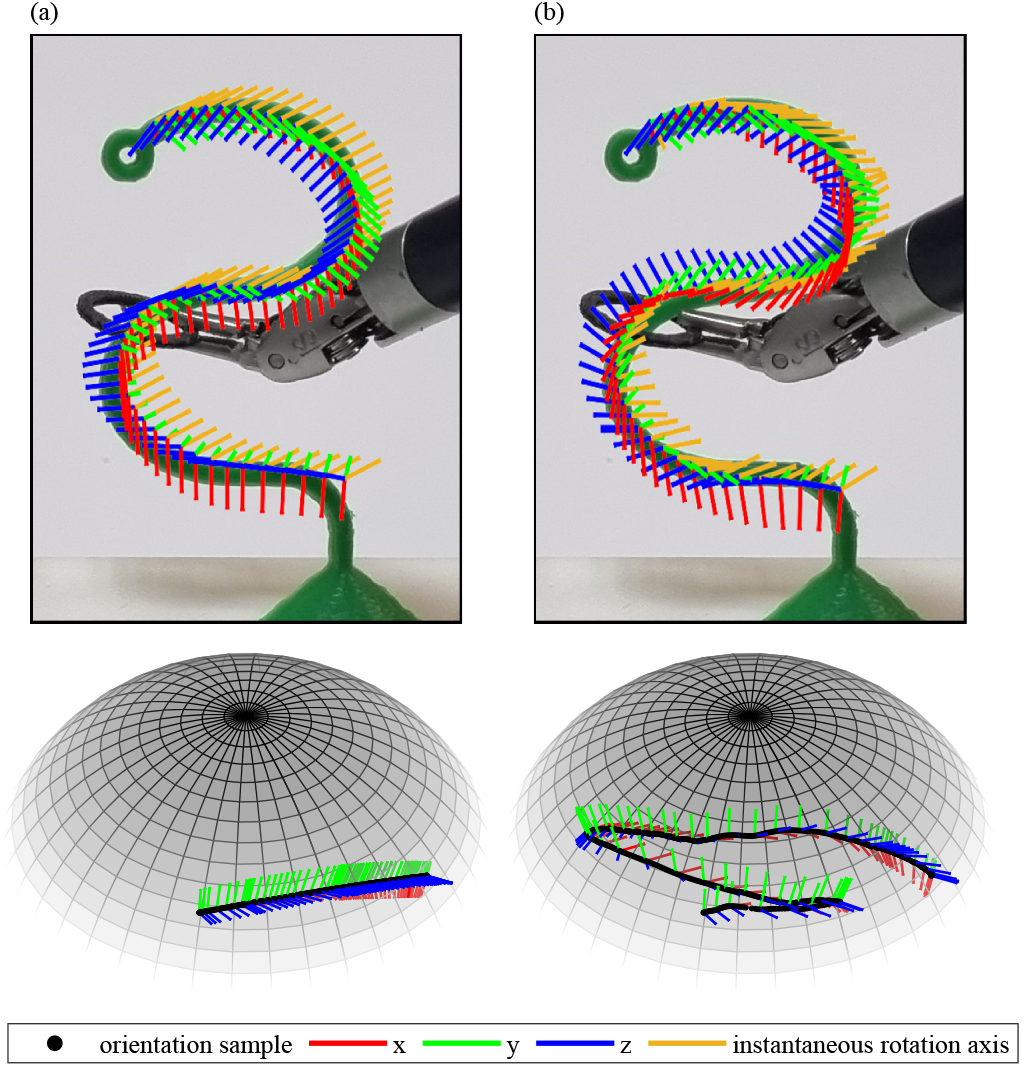
Analysis of angular speed and local geometry of the orientation path of a teleoperated robot-assisted surgical instrument’s tip. The top row presents the transfer of a ring on a wire, and the red, green, and blue lines represent a moving frame attached to the rotating gripper of the tool-tip. The orange line represents the instantaneous rotation axis (represented in a fixed reference frame) for the transition between two subsequent frames. The spheres in the bottom row illustrate the same movements as samples of orientations (black dots) using quaternions in 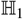 (for intuition, we illustrate 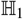 on which unit quaternions lie using the 2-sphere). The angular speed is determined by the spherical distance between consecutive orientations in 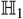, and the local curvature is determined by the curvature of the path’s projection in 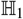 onto the tangential plane (see Methods). (a) The tool-tip rotates around a fixed instantaneous rotation axis, and therefore, it creates a geodesic in 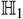 (arc). (b) The instantaneous rotation axis varies, therefore, the orientation path is a non-geodesic.

To discover the building blocks of human movement control, experimental and computational approaches are used. For example, the hand is kinematically redundant, and the basis on which the nervous system chooses one trajectory (path and timing) from an infinite number of possibilities has been thoroughly studied [4], [5], [6]. Motor invariants, sometimes also referred to as motor primitives, are robust patterns that can be quantified in human movements. They are typically observed across repetitions of the same movements within and between participants, and are considered to be a result of active control by the nervous system [4].

One such invariant, which may result from the optimization of movement smoothness [4], is the straight path and bell-shaped velocity trajectory that characterizes fast point-to-point movements. Another prominent invariant, which has also been linked to movement smoothness [7], is the speed-curvature power law [8], which describes a relation between the instantaneous linear speed (*ν*) of a two-dimensional translational movement and its local path curvature (*κ*):

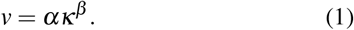

The linear speed characterizes the movement in space and time, whereas the curvature is a pure geometrical property – the local deviation of the path from a straight line. The speed gain (*α*) is generally piece-wise constant [8], [9], and the exponent (*β*) is negative and constant for similar paths.

This law was first reported in the case of drawing ellipses, with an exponent value of −1/3 [8], and was later generalized to a spectrum of power laws whose exponent depends on the number of curvature oscillations per angle unit [10]. It has been demonstrated in a variety of movements, including movements of the hand [8], feet [11], and even the head during walking [12]. The law is thought to rely on physiological mechanisms, but its origin remains controversial. Some studies argue that the trajectory is planned within the central nervous system [7], [13], [14], [15]. Others attribute the law to lower levels of the sensorimotor system [16], or to the biomechanics of the limbs [17]. The neural representation of the power law was also argued to affect perception – humans perceived animated motion as more natural when it obeyed the speed-curvature power law [18]. A similar representation may underlie the recent results suggesting that human-robot physical interaction is most efficient when the robot moves in accordance with the speed-curvature power law [19].

When movements in three dimensions were considered, the power law was extended to the speed-curvature-torsion power law [20], which accounts for deviation from a straight and a planar path. This extended power law was demonstrated in surgical suturing, and shown to be dependent on the level of user expertise, and on whether the task was performed via RAMIS or via open surgery techniques [21]. However, this is still not sufficient to account for fine manipulation in many real-life robotic applications, such as RAMIS [22], that involve both translation and rotation that may be independently controlled [23]. Indeed, while motor invariants are extensively studied in the control of translation, a similar understanding of the control of rotation is lacking.

Some studies have already started to fill the gap. For example, a study of planar path following revealed that users exploited more portions of arm redundancy when the task required the control of both position and orientation of the tool compared to position alone [24]. Similarly, in the context of RAMIS training, peg transfer with a change of orientation [25], ring-on-wire transfer [26], and needle driving [27], [28], were investigated. However, a fundamental computational understanding of human control of orientation, and subsequently, manipulation in six degrees of freedom, has yet to be developed.

In this study, we aimed to address this gap by quantitatively characterizing the relation between the spatio-temporal and spatial changes of orientation trajectories represented by unit quaternions in 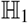 (Fig. 1). We propose a new power law linking the speed to the geometry in human control of movement orientation. We demonstrate this law by analyzing movements of surgical residents performing a ring tower transfer using the da Vinci Research Kit (dVRK, Fig. 2a-b), a RAMIS research platform [29]. This task is commonly used in training of robotic surgeons [30], and requires the manipulation of the position and the orientation of a ring held by the gripper of the teleoperated instrument. We also demonstrate this law on a needle insertion task teleoperated by surgeons using the da Vinci Surgical System (dVSS, Intuitive Surgical, Inc., Sunnyvale, CA) taken from the publicly-available JIGSAWS dataset [31].

**Fig. 2.**
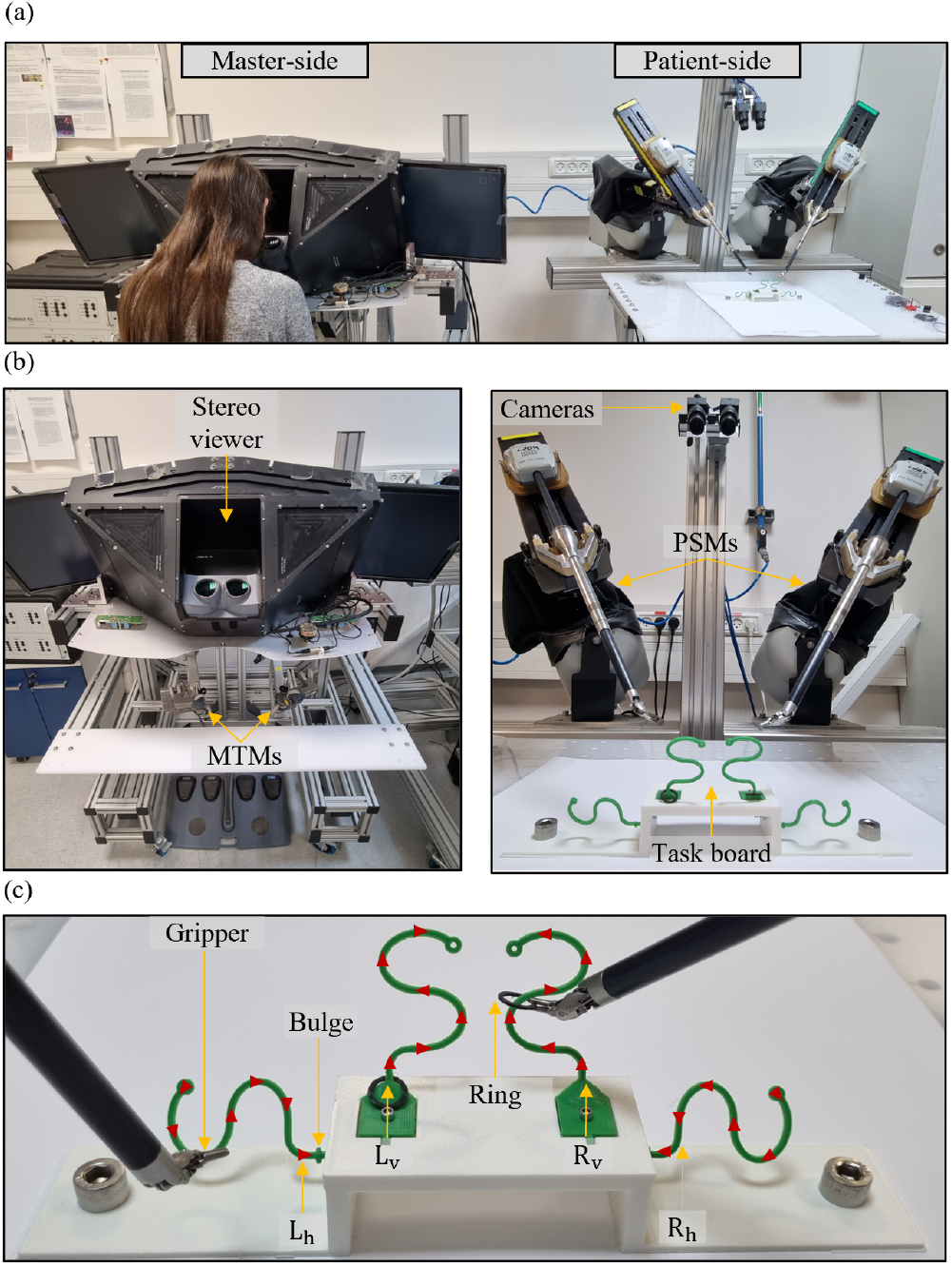
Experimental setup and task. (a) The dVRK overview with master and patient sides. (b) The master side consists of a stereo 3D viewer and MTMs. The patient side consists of cameras, PSMs, and a task board. (c) The task board with red arrows depicting the instructed ring paths.

The remainder of this paper is organized as follows. In Section II we present the mathematical representation of orientation, the definition of the power law, and the description of the data and the analysis. In Section III, we present the results of fitting the power law to the data, and validation studies. In Section IV, we discuss the results in relation to the translational power law and their implications in the context of RAMIS, the limitations of this study, and future studies. In Section V, we present our conclusions.

## II. METHODS

### A. Mathematical formulation

In this subsection, we present the formulation of a new power law which links the speed to the geometry of the orientation of teleoperated robotic instruments. The manipulation of a rigid body in three-dimensional space involves movement in six degrees of freedom. Three degrees of freedom are associated with translation – the movement of the origin of a reference frame that is attached to the body, and the other three are associated with rotation of the reference frame about some axis. We considered rotations represented by unit quaternions, denoted by 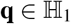. 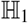 is the space of unit quaternions and is also known as the 3-sphere – a four-dimensional unit size sphere. This is helpful when visualizing quaternions (Fig. 1, bottom row).

Quaternions are hyper-complex numbers comprised of a real part 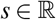 and an imaginary part 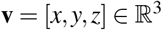, such that **q** = *s* + **i**x + **j**y + **k**z, **i**^2^ = **j**^2^ = **k**^2^ = **ijk** = −1, **ij** = **k**, **ji** = –**k**. According to Euler’s theorem, for any 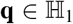, there exists an angle *θ* ∈(–*π, π*] and an axis 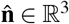, such that:

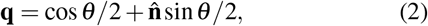

where **q** is the quaternion that rotates a point in 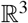 by *θ* around 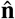.

Let 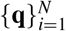 be a discrete time *N* samples orientation trajectory (e.g., of the tool-tip of the Patient Side Manipulators (PSMs) of a teleoperated surgical robot – see next section for details), such that 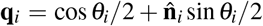 is the quaternion that rotates all the vectors represented in a fixed frame (**q_I_** = 1) by an angle *θ_i_* around an axis 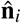. We denote the discrete time trajectory of the transition quaternion as
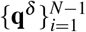, such that 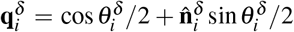 is the quaternion that rotates **q**_*i*_ to **q**_*i*+1_ with an instantaneous angle 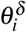 around an instantaneous axis 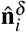:

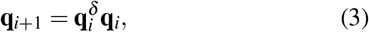

where *δ* is used to indicate an incremental and small transition between **q**_*i*_ and **q**_*i*+1_. The angular speed at sample *i* (*ω*_i_) quantifies the spatio-temporal change of the PSM tool-tip’s orientation when it rotates about its center of mass:

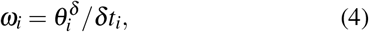

where *δt*_*i*_ = *t*_*i*+1_ – *t_i_* is the time sampling interval. The curvature at sample *i* (*λ_i_*) quantifies the spatial change of the instantaneous axis of rotation as the local curvature of a curve 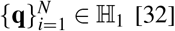 [32]:

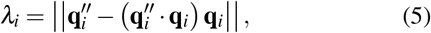

where ||·|| is the 2-norm. We approximate the second derivative of a quaternion curve at sample *i* as in [32], [33]:

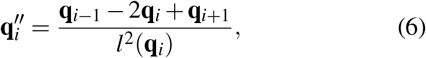

where *l*(**q**_*i*_) = cos^−1^(**q**_*i*−1_ · **q**_*i*_)/2 +cos^−1^(**q**_*i*_ ·**q**_*i*+1_)/2. Eq. 5 originates from the projection of a curve in 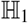 onto the tangential plane, so it promises that a geodesic in 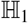 is characterized with *λ* = 0 along the great arc.

The proposed power law relates the angular speed of the tool-tip with the local curvature of its path:

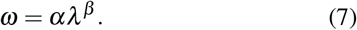

### B. Datasets

To demonstrate the proposed power law in a natural orientation control task, we analyzed recordings of a teleoperated bi-manual ring tower transfer task (Fig. 2c) performed with the dVRK (Fig. 2a-b). Specifically, we analyzed the kinematics of the patient side manipulators (PSMs) that were used to perform this task. These data are part of a large dataset that was collected at the Biomedical Robotics Lab at Ben-Gurion University of the Negev as part of a long-term surgical skill learning study.

18 surgical residents (16 males and 2 females, aged 27-40, 17 right-handed and 1 left-handed) from the Soroka Medical Center participated in the experiment after signing an informed consent form approved by the Human Participants Research Committee of Ben-Gurion University of the Negev, Beer Sheva, Israel. None of the participants had prior RAMIS experience. The participants were seated in front of a surgeon’s console of the dVRK [34] and used the Master Tool Manipulators (MTMs) to control two PSMs (Fig. 2a). Each participant took part in four sessions of the experiment; each session was comprised of three trials that occurred before, during, and after a hospital shift. The gap between two consecutive sessions was near one month.

Each trial consisted of four ring tower transfers (Fig. 2c) – two vertical towers (R_v_,L_v_) and two horizontal towers (R_h_,L_h_) relative to the task board (see Video 1 for a demonstration of the task). At the beginning of each trial, the participants grasped the ring at the bottom of tower R_v_ using the right PSM, and were instructed to trace the wire without touching it and extract the ring at the top of the tower. Following that, participants transferred the ring to the left PSM, inserted it through tower L_h_ and placed it after the bulge. Next, participants used the left PSM to grasp and extract a second ring from tower L_v_ in a similar manner, and then transferred it to the right PSM and inserted it through tower R_h_. All the participants watched an instructional video explaining the task and showing one successful performance. Participants were instructed to avoid touching the wire, ring drops, PSMs collisions with the task board and between themselves, and to complete the task as fast as possible.

If the power law is a robust motor invariant, it is expected to be valid for different tasks and teleoperation systems. Hence, we analyzed another dataset – a subset of the JIGSAWS dataset [31], which contains the kinematic and video data of teleoperated surgical training tasks performed with the dVSS by eight surgeons with varying experience levels. Our law describes movements with a considerable change of orientation. Hence, we analyzed only the kinematics of the right PSM in the right-hand needle insertion segments from the suturing trials (a total of 164 segments).

### C. Data analysis

We recorded the ring transfer orientation trajectories of the tool-tips of both PSMs as quaternions at 100 [Hz], and the video at 35 [Hz]. We excluded from the analysis 21 out of 864 segments, in cases of: robot shut down due to joint limits or excessive forces, dropped ring by participant, or the use of incorrect PSM (2.43%, 1 R_v_, 4 R_h_, 6 L_v_, and 10 L_h_). Participants failed more in the left hand towers, likely due to their right hand dominance, and in the horizontal towers, due to the proximity to the edge of the workspace.

We analyzed the data with a custom-written MATLAB code (The MathWorks, Natick, MA, USA). In the ring transfer task, each trial was manually segmented into the four ring tower transfers using the video of the trial. The transfers at towers R_v_ and L_v_ began when the PSM’s jaw grasped the ring and ended when the ring was extracted. The transfers at towers R_h_ and L_h_ began when the ring was inserted into the tower and ended when it was placed after the bulge.

In both datasets, we low-pass filtered the orientations using spherical linear interpolation [32]. Then, for each sample we calculated the angular speed (*ω_i_*) and the local curvature (*λ_i_*). Next, for all the samples in each segment, we fit a log-log linear regression model:

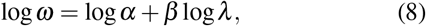

where log is the natural logarithm. To assess the goodness of fit of the proposed power law, for each segment, we calculated the variance explained by the linear model of the log-log transformed data (R^2^), and evaluated its mean and standard deviation over all the segments. To asses the variability of the model parameters and explained variance, we used the coefficient of variation (the standard deviation divided by the absolute mean). To show that the law’s parameters are consistent throughout all repetitions of the ring transfer task, we calculated the bootstrap 95% confidence interval (CI) for the means of *α* and *β* in each session, trial, and tower.

### D. Validation studies

We made several efforts to ensure that our proposed power law is not a consequence of the way we estimate the angular speed and curvature from the ring transfer dataset. First, to verify that the model parameters are not sensitive to the sampling rate of the orientation trajectory, we downsampled the data to 50 [Hz] and 33 [Hz], and compared the distributions of the fit parameters and the explained variance.

Next, we manipulated a recorded orientation trajectory by sampling the quaternions at close to equal arc-lengths in H_1_. By assigning the original timestamps to the manipulated data, we generated a trajectory with a similar path, yet an angular speed that violates the power law. In addition, we simulated an orientation trajectory (not based on the datasets) that is inconsistent with the power law – it has two curvature oscillations and a nearly constant angular speed. We then fit the power law parameters to these manipulated and simulated examples of data, and verified that in these cases that violate the power law, we indeed do not find a good fit.

## III. RESULTS

### A. Power law analysis

Fig. 3 presents an example of the analysis of one ring tower transfer. In analyses of speed-curvature power laws, it is important not to fit the power law to geodesics (i.e., straight paths in the case of translation or fixed instantaneous rotation axes in the case of orientation) [35]. A geodesic in 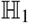 (Fig. 1a) results in an arc in the path of the quaternion axis 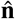 (Eq. 2). The example path in Fig. 3a (top sphere) clearly deviates from an arc, implying the existence of a spatial change of orientation. Accordingly, the dispersed instantaneous rotation axis path in the bottom sphere indicates a change in the direction of rotation. A large spherical distance between consecutive instantaneous rotation axes implies a large curvature, and vice versa. The log-transformed curvature values in the ring transfer data ranged from log *λ* = −2.06 to log *λ* = 15.42 (log *λ* = 4.73 ± 1.86, mean ± standard deviation), and from log *λ* = −1.34 to log *λ* = 14.42 (log *λ* = 4.43 ± 2.13) in the needle insertion data. The large range of values indicates that we do not fit the model to geodesics (i.e., *λ* > 0).

**Fig. 3.**
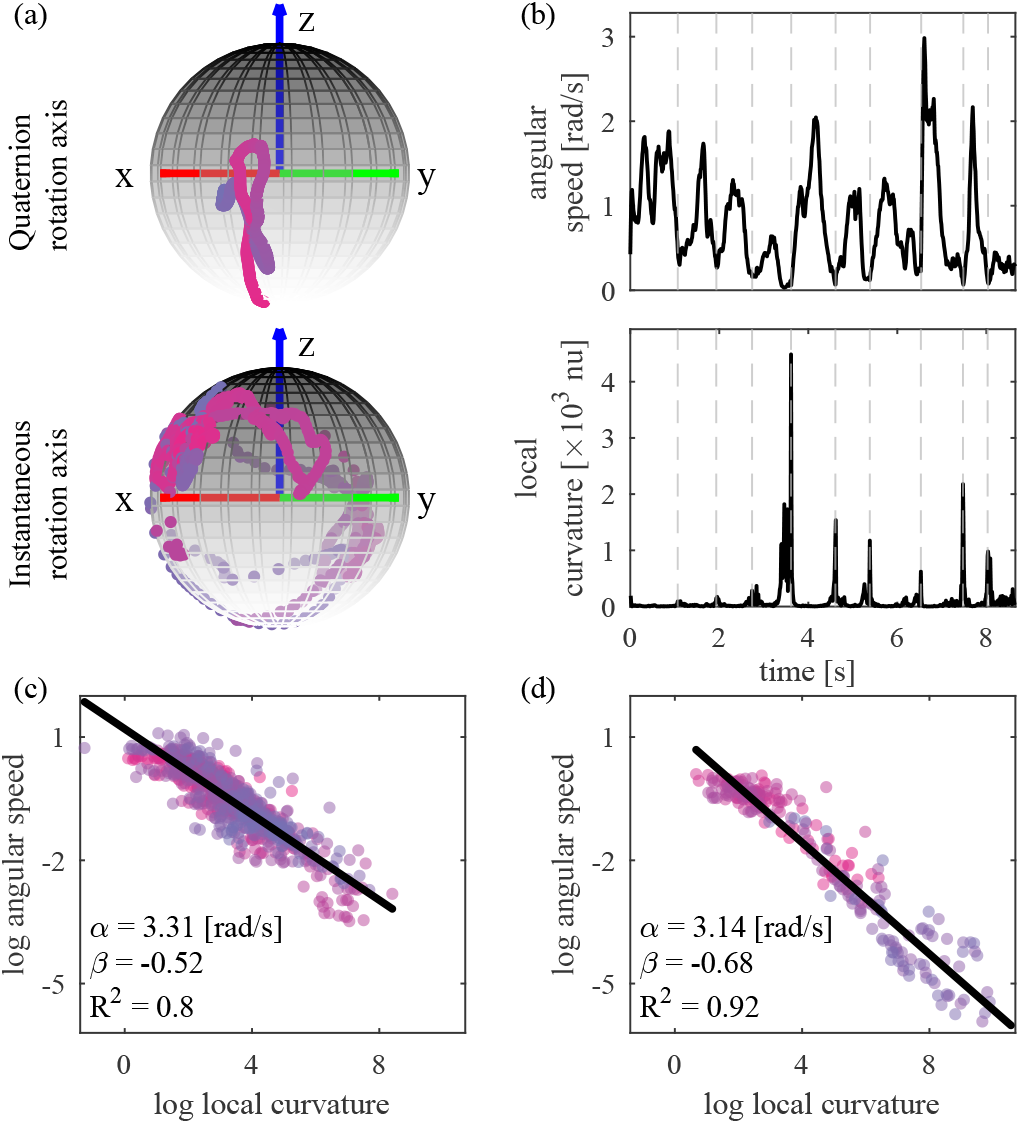
Examples of power law analyses. (a) An extrinsic reference frame is shown as red, green, and blue arrows. The color of the points changes from pink (beginning) to purple (end) to indicate time progression. In the ring transfer, the tool-tip did not follow a geodesic in 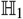. This is observed in the quaternion rotation axis path (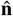, top sphere) that deviates from an arc, and in the instantaneous rotation axis path (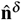, bottom sphere) that is widely dispersed. (b) Angular speed (top) and local curvature (bottom) of the tool-tip orientation trajectory in the ring transfer. The dashed lines indicate peaks in the local curvature that correspond to valleys in the angular speed. nu - no units. The log-log regression models (black lines) are fit to the data (dots) in the (c) ring transfer task and the (d) needle insertion task.

Examining the angular speed and curvature trajectories in Fig. 3b clearly reveals that the angular speed is low when the local curvature is high, and vice versa. This is additionally supported by the log-log regression that is depicted in Fig. 3c, which presents a speed gain *α* = 3.31 [rad/s], an exponent *β* = −0.52 and a good fit to the data with R^2^ = 0.8. The distributions of the fit parameters to the ring transfer data sampled at 100 [Hz] and the explained variance from all the segments (including different participants, sessions, trials, and tower segments) are depicted in Fig. 4a. Both the speed gain and the exponent are narrowly distributed around the mean (the coefficient of variation was 0.23 for the speed gain and 0.09 for the exponent). In addition, the model resulted in a consistently high explained variance, which was also narrowly distributed (the coefficient of variation was 0.06). The explained variance was similar to the values that were previously reported in studies of the speed-curvature power law in translational movements [20], [36]. This analysis suggests that, consistently with the proposed power law in Eq. 7, the angular speed was exponentially related to the local curvature with a speed gain of *α* = 3.39 ± 0.78 [rad/s], and an exponent of *β* = −0.54 ± 0.05. The variance explained by the model was R^2^=0.81 ±0.05.

**Fig. 4.**
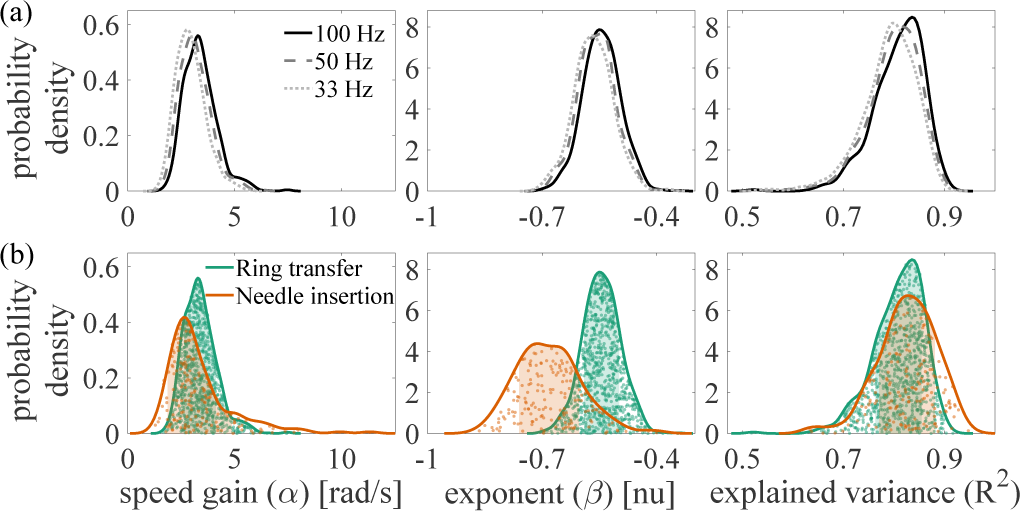
The distributions of the fit model parameters. (a) The distribution in the ring transfer task, and the effect of sampling rate. (b) The distributions in the ring transfer task (green), and the JIGSAWS needle insertion task (orange). The scatter plots show the model parameters and explained variance. The shaded areas mark the mean±standard deviation.

Our analysis of the needle insertion segments in the JIGSAWS dataset further supports the robustness of our proposed power law. An example of a log-log model of one needle insertion from the JIGSAWS dataset is presented in Fig. 3d, and Fig. 4b shows the distributions of the fit parameters in the needle insertion task compared to the ring transfer task. The parameters are also narrowly distributed (the coefficient of variation was 0.47 for the speed gain, 0.12 for the exponent, and 0.07 for the explained variance). Yet, the speed gain is slightly more variable (*α* = 3.37 ± 1.59 [rad/s]) compared to the ring transfer task, and the exponent is lower and more variable (*β* = −0.68 ±0.08). The explained variance (*R*^2^ = 0.82 ±0.06) is similar in both tasks.

In addition to demonstrating the adherence of rotational movements to the new power law, in the ring transfer task, we assessed whether the parameters of the power law are consistent across different sessions, trials and tower segments (Fig. 5). Since this is the first study to report the power law, we did not support this analysis by a statistical model, but rather present the distributions, their means and bootstrap 95% CIs. The similarities between CIs suggest that the model’s parameters were robust with respect to these factors. With the progression of the sessions, the speed gain seems to decrease and the exponent appears to increase, but the change is very small (an average decrease per session of 0.07 [rad/s] in the speed gain, and an average increase per session of 0.01 in the exponent). In addition, the speed gains in the horizontal towers segments were slightly larger than in the vertical towers segments. This could be due to slower rotations near the edge of the workspace in the horizontal towers. We also found slightly larger exponents in segments performed with the right PSM compared to the left PSM, which may be attributed to the right-handedness of our surgeon participants, or to a difference in sensorimotor control of right and left hand rotations. However, these variations may also stem from the fixed order of segments in each trial.

**Fig. 5.**
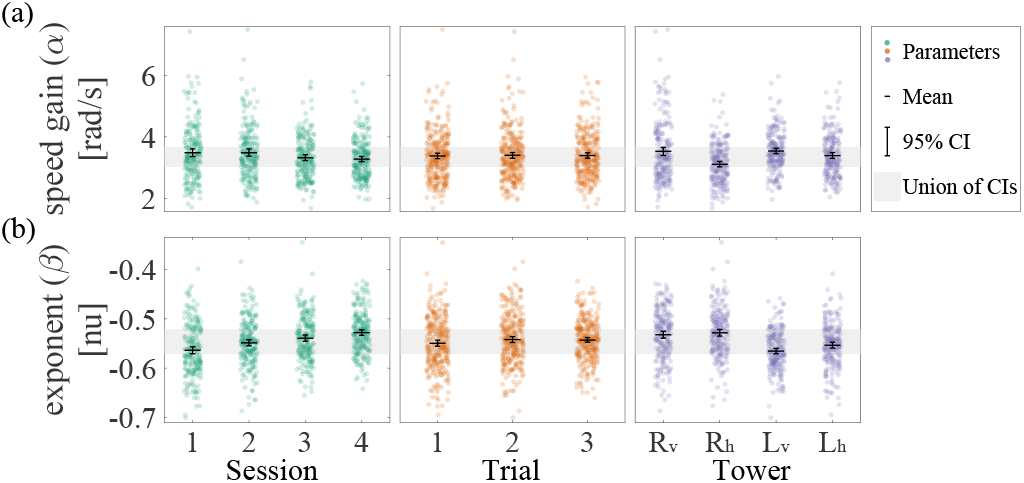
Summary of the power law parameters in the ring transfer task. The horizontal lines show the mean of the (a) speed gain and (b) exponent in each session, trial and tower segment (left, middle and right panels). The error bars indicate the 95% bootstrap CI for the mean. The gray shaded areas mark the union of all CIs for each model parameter.

### B. Validation studies

One possible criticism to the validity of the proposed power law is that it could be a result of the sampling process [37]. If this were the case, we would expect the parameters of the power law to depend on the sampling rate. Hence, in the ring transfer task, we downsampled the data to 50 [Hz] and 33 [Hz] and repeated the power law analysis. Fig. 4a shows the distributions of the fit parameters in these two cases compared to those of the original sampled data. We observed a slight shift in the distribution of the speed gain with sampling rate (50 [Hz]: 3.16±0.74, 33 [Hz]: 2.94±0.69), and a nearly identical distribution of the exponent (50 [Hz]: −0.55 ±0.05, 33 [Hz]: −0.56 ±0.05) and of the explained variance (50 [Hz]: 0.8 ±0.05, 33 [Hz]: 0.79±0.05).

Another possible criticism is that the results are purely a computational consequence of the math used to compute the angular speed and local curvature [38]. If this were the case, any dataset that we would generate would yield a good fit of the power law, regardless of whether the actual trajectory and path obey it. To counter this argument, we manipulated an existing ring transfer orientation trajectory by breaking the dependency between the speed and the geometry. Fig. 6a shows the results of the manipulated orientation trajectory, and the poor fit of the power law. In addition, we simulated an equally spaced orientation trajectory in 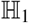, thus creating a path with a varying curvature traveled in a constant angular speed (Fig. 6b). The generated path is equivalent to an ellipse in an Euclidean space, which has two curvature oscillations, and similarly does not yield a good fit to the power law.

**Fig. 6.**
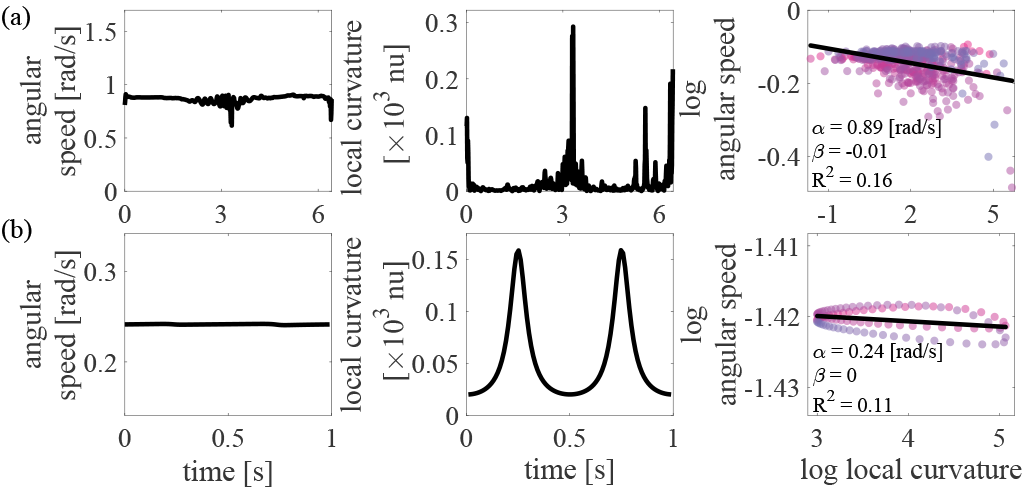
Manipulated and simulated orientation trajectories that violate the power law. (a) We manipulated a ring transfer orientation trajectory by sampling at close to equal arc-lengths resulting in a nearly constant angular speed (left) and curved path (middle) which violates the power law (right). (b) We simulated an orientation trajectory with nearly constant angular speed (left) and two curvature oscillations (middle) which is inconsistent with the power law (right). The color change is as in Fig. 3.

## IV. DISCUSSION

We presented a new power law which links the angular speed to the geometry of the tool-tip orientation in teleoperation of a robot-assisted surgical system. We analyzed kinematic data recorded during teleoperated surgical training tasks – transfer of a ring along a wire and needle insertion. Unlike most motor control studies, we focused on the orientation of the controlled object, rather than on its position. We addressed the hypothesis of a power law linking between the local spatio-temporal change of orientation – the angular speed of the PSM’s tool-tip when it rotates about its center of mass, and the local spatial change of orientation – the way a curve in 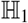 bends with respect to a geodesic. The model successfully accounted for 81% of the variance in the variables which spanned over at least two orders of magnitude, making the model a legitimate power law [39].

When formulating our new power law we were inspired by the well-known speed-curvature law describing translational movements [8]. Our proposed law extends the original law to account for rotational movements. Indeed, the variance explained by our power law does not fall short of that found in translational 3D movements in the general [20], [36] as well as in the surgical context [21]. Similar to the original power law, our results suggest the existence of an underlying control process that dictates dynamic commands in response to geometrical input. However, the level of the sensorimotor system in which this process is implemented, as well as which control methods are utilized, remain unknown. Furthermore, it remains to discover in future work whether in general pose control tasks, the translation and rotation are controlled separately (as was suggested for reaching movements [40]), or simultaneously [6], as well as whether linear and angular speed are affected in more complex ways by the local geometries of translational and rotational paths.

The parameters of the power law were narrowly distributed across participants and repetitions of the ring transfer task and the needle insertion task from the JIGSAWS dataset. This suggests that the power law is a robust motor invariant of the control of orientation. The distribution of the exponent in the needle insertion was more negative and wider compared to the ring transfer, which may imply the existence of a task-dependent family of power laws, as in the translational law [10], [20]. The variability could also stem from the needle insertion task being less structured than the ring transfer task. These hypotheses may be tested in the future using wire designs with diverse curvature profiles.

The good fit of the data to our proposed model does not indicate that it is necessarily true. Despite the fact that we did not constrain the tool to move according to any specific velocity profile, we considered the option that the results are either sampling-related or a trivial mathematical consequence [38]. Downsampling the orientation trajectory caused a small shift in the distribution of the speed gain, and nearly no change in the distribution of the exponent and explained variance. The relatively larger sensitivity of the speed gain to the sampling compared to the exponent and the explained variance may be explained by the relatively larger dispersion of the speed gain compared to the exponent and the explained variance also in the original fit. Therefore, we concluded that the power law model is insensitive to the sampling rate. Additionally, we manipulated the data to violate the power law, and simulated data inconsistent with the power law. Both yielded a poor fit to the law, thus rejecting its triviality. Nevertheless, our lack of ability to refute the model does not mean it is true, as “all models are wrong, but some are useful” [41]. We nevertheless believe that our proposed power law has important implications in the context of RAMIS, as we outline in the next paragraphs.

The parameters of the translational speed-curvature power law were previously proposed as a surgical skill signature [21], [42]. Our goal in this study was to establish the power law, and hence, we did not make any predictions about how its parameters may reflect skill. Nevertheless, posthoc analysis revealed only slight changes in the parameters over the sessions. However, the data spanned only four training sessions over four months, and the timescale for learning complex rotations has not yet been established. Hence, to link the parameters of the power law to surgical skill, testing movements of a large sample of surgeons of various expertise levels, or years of monitoring surgical skills, are needed.

Similar to the speed-curvature power law in translations, the new power law suffers from singularity at *λ* = 0, as the angular speed should become infinite. Though we did not force participants to rotate around a varying instantaneous axis, our analysis showed that they did. Nevertheless, low curvature points were observed. Future work is needed to construct an analytic framework that explains the new power law and predicts its extension to geodesics, akin to the minimum-jerk framework in the control of translation [7]. This would facilitate understanding of rotation coordination across a wider range of local curvatures. In drawing, for example, points of inflection are used to segment movements [9] and it remains to be determined if the segmented control hypothesis also holds for the control of rotation.

We used teleoperated RAMIS platforms to study the control of the orientation of a remote tool. The dynamics of any apparatus or environment constrain the human behavior: tool manipulation in thin air is constrained by the inertia and the friction of the tool, and to perform movements in viscous environments users adapt to the dynamics of the environment [43]. Teleoperation influences the smoothness of the movement [21], [44], and the design of the teleoperation system can influence the empirically derived parameters of the power law. However, so would any tool or manipulated object. Moreover, the structural significance of the power law lies in the robustness of the fit parameters across different repetitions of the same task by different participants, regardless of the experimental platform. Finally, some designs could be so unnatural that they would cause even fundamental motor invariants to breakdown, as shown previously for the case of adherence to psychophysical laws [45], [46]. Hence, in future studies, the goodness of fit of our new power law could be incorporated in benchmarking of the transparency of teleoperated systems.

Models of human motor control can be harnessed to a wide range of applications in human-centered control of robotic systems. Such systems may implement virtual fixtures – software generated dynamic signals applied to the user [47]. In RAMIS, virtual fixtures can assist surgeons when navigating around delicate anatomical structures [48], [49]. A study of translational movements showed that such robotic assistance is most effective if it obeys laws of human movement, as users applied less force on the robot when the guided movement complied with the speed-curvature power law [19]. The new power law can be used to design guidance virtual fixtures in tasks with rotational movements by adding constraints to the motion planning algorithm. Surgeons may be guided by adjusting the virtual fixtures to each surgeon’s unique parameters, or to the parameters of expert surgeons.

Automation of surgical procedures, such as the recently reported autonomous small bowel anastomosis [50], is another promising research direction that can provide better consistency across operations. While full surgical procedures are not yet automated, collaboration between surgeons and autonomous assistants is expected in the near future. Using biologically plausible movement in robots makes their motion predictable and increases the efficiency of human-robot cooperation [1], [2], [3]. Therefore, we speculate that surgeons would be more comfortable operating in collaboration with robots that move in accordance with our proposed power law. The new power law could be combined with the power law for translational movements [21], [42], and yield trajectories that could account for all six degrees of freedom in space, and instruct rate according to the power laws.

However, it is important to note that clinical procedures involve more complex movements than what we studied here. They also involve time-dependant perturbations, such as the movement of the beating heart, and visual remapping due to tool or camera viewpoint misalignment [51], all of which could affect task performance [52], [53] and surgical outcome. To design virtual fixtures and autonomous surgical assistance with the help of motor invariants such as the power law that we report here, future studies are needed to test the effect of dynamic and visual perturbations on the parameters of the power law, and its applicability to more complex movements in contact with tissue.

## V. CONCLUSIONS

We reported a novel power law describing the relation between the local geometry of the orientation path of a remote controlled tool and the angular speed used to follow that path. The power law analysis yielded a good fit of the experimental data to the model, with a consistent speed gain and exponent across participants and repetitions of the ring transfer and needle insertion tasks. The results suggest the existence of a controller that translates the local spatial characteristics of the orientation of a rigid body into executable motor commands. We argue that human control of orientation should be studied thoroughly and can be applied to the design of human-centered teleoperation systems in many applications requiring fine manipulation, such as RAMIS.

## ACKNOWLEDGMENT

We thank Tami Matus for the administrative assistance and Noa Yamin, Sapir Goldring, Netali Auerbach, Nadav Amitai, Yuval Kassif and Ehud Zippin for collecting the data.

