## Supplementary material for "A New Power Law Linking the Speed to the Geometry of Tool-Tip Orientation in Teleoperation of a Robot-Assisted Surgical System": Video 1: README.txt

The supplementary material include one video with recording of a single trial, including four ring tower transfer segments.

Video 1 can be played using Windows Media Player, version 12.

Contact information: Or Zruya
