## Supplementary material for "A New Power Law Linking the Speed to the Geometry of Tool-Tip Orientation in Teleoperation of a Robot-Assisted Surgical System": Video 1: Summary.txt

The multimedia materials attached to this submission is one video file.

Video 1 demonstrates a single trial of the experiment, which consists of four ring tower segments.

An annotation is presented describing:
1 - The manipulator that is currently used (right or left)
2 - The orientation of the tower with repsect to the white task board (vertical or horizontal). 
